# A local insulin reservoir ensures developmental progression in condition of nutrient shortage in *Drosophila*

**DOI:** 10.1101/2021.07.05.451134

**Authors:** Suhrid Ghosh, Weihua Leng, Michaela Wilsch-Bräuninger, Pierre Léopold, Suzanne Eaton

## Abstract

Insulin/IGF signalling (IIS) controls many aspects of development and physiology. In *Drosophila*, a conserved family of insulin-like peptides (Ilp) is produced by brain neurosecretory cells and exerts systemic functions. Here, we describe the local uptake and storage of Ilps in the Corpora Cardiaca (CC), a group of alpha cell homolog that produces the glucagon-like hormone AKH. Dilp uptake relies on the expression of Impl2, an IGF-BP that accumulates in the CCs. During nutrient shortage, this specific reserve of Ilps is released and activates IIS in a paracrine manner in the prothoracic gland, securing accelerated entry into pupal development through the production of the steroid hormone ecdysone. We therefore uncover a sparing mechanism whereby local Ilp storage and release activates the production of steroids and ensures early developmental progression in adverse food conditions.

**Highlights:**

- Dilps are uptaken by CC cells through the IGF-BP Imp-L2
- the CC-Dilp store is released upon nutrient shortage and activates IIS through CC projections on the PG
- upon nutrient shortage, IIS activation in the PG ensures an accelerated transition from larval feeding stage to metamorphosis.

## Introduction

Insulin and insulin-like growth factors (IGFs) are conserved modulators of growth and metabolic homeostasis. They are produced by specific endocrine organs in response to nutrient availability and stimulate peripheral tissues through two main routes. Circulating insulins and IGFs act in an endocrine manner on distant organs through mechanisms of secretion and action that have been extensively studied (Campbell & Newgard, 2021; Petersen & Shulman, 2018; Tokarz, MacDonald, & Klip, 2018). In several instances, insulin/IGFs are also produced and act locally. This is the case for IGF-I produced in peripheral organs like bone chondrocytes, which acts in an autocrine manner (Wang, Bikle, & Chang, 2013). Local insulin/IGF signalling is also key for the development and homeostasis of the central nervous system, which is separated from the systemic circulation by the blood-brain barrier (Fernandez & Torres-Alemán, 2012). In mammalian brains, IGF-I expression is spatially and temporally controlled, participating in the rapid proliferation of neural stem cells (NSCs) (Popken et al., 2004). IGF-I is expressed in neurons, astrocytes and NSCs, contributing to both paracrine and autocrine signaling (D’Ercole & Ye, 2008).

In the *Drosophila* model, a subset of insulin/IGFs called *Drosophila* insulin-like peptides (Dilps) is expressed in neurosecretory cells and contributes to systemic activation of insulin/IGF signaling (IIS). Similar to mammals, local brain expression of Dilps in glial cells also contributes to coupling NSC proliferation with systemic metabolic status (Yuan, Sipe, Suzawa, Bland, Siegrist & Id, 2020; Spéder & Brand, 2018).

In parallel, local activation of IIS can be triggered by alternative routes, providing the possibility to uncouple insulin/IGF signaling from nutrient availability. The role of this uncoupling is exemplified in the case of “brain sparing” a phenomenon observed in many species, including human, whereby moderate nutrient deprivation affects primarily body growth, while brain growth is preserved (Cox & Marton, 2009). The molecular mechanism of brain sparing has been addressed in *Drosophila*. It is associated with the specific activation of neuroblast growth through local production of the ligand Jelly belly (Jeb) and activation of the Anaplastic lymphoma kinase (Alk) receptor upstream of PI-3-kinase, even in the absence of nutrients (Cheng et al., 2011).

Here, we unravel a distinct mechanism locally uncoupling insulin release from the presence of nutrients and leading to the maintenance of IIS activation in a key endocrine organ during development. The initial observation of a pool of Dilps accumulating in an endocrine gland called the corpora cardiaca (CC) (S. K. Kim & Rulifson, 2004), suggested the possibility that these insulins could activate IIS under specific conditions, in a way distinct from their systemic function. Indeed, the CC cells do not produce insulins, but rather a glucagon-like hormone called AKH, whose release is stimulated by nutrient shortage. Our present work unravels some features of the mechanism of uptake of insulins by the CCs. We show that, conversely to systemic insulins, CC insulins are released in response to nutrient deprivation. We further demonstrate that the local release of CC insulins upon starvation maintains high IIS levels in the prothoracic gland (PG), which produces the steroid ecdysone. Finally, we establish that the local activation of IIS in the PG secures a fast transition to pupal development in absence of nutrients.

Therefore, our work uncovers a distinct mechanism of organ-sparing, whereby a pre-accumulated store of Dilps is released in adverse food conditions and ensures fast developmental progression through steroid production.

## Results

### Corpora cardiaca cells take up Dilp2 and 5

The Corpora Cardiaca (CC) is part of the ring gland (RG), a composite endocrine organ located in the vicinity of the larval central nervous system (CNS) (Figure 1a). Apart from the CC, two other endocrine tissues - the prothoracic gland (PG), which synthesizes ecdysone, and the corpora allata (CA), which synthesizes the juvenile hormone – together form the PG (Figure 1b). CC cells are neurosecretory in nature and extend processes that terminate in the larval aorta and the PG (Figure 1b). The IPCs, located in the pars intercerebralis of the larval brain, also project to the aorta (Figure 1b). Previous work describes a strong accumulation of anti-Dilp2 immunoreactivity in the corpora cardiaca (CC) (S. K. Kim & Rulifson, 2004), in addition to the one observed in the cell bodies of IPCs (Rulifson, 2002). However, whether CC cells express the Dilp2 gene has not been investigated.

**Figure 1.**
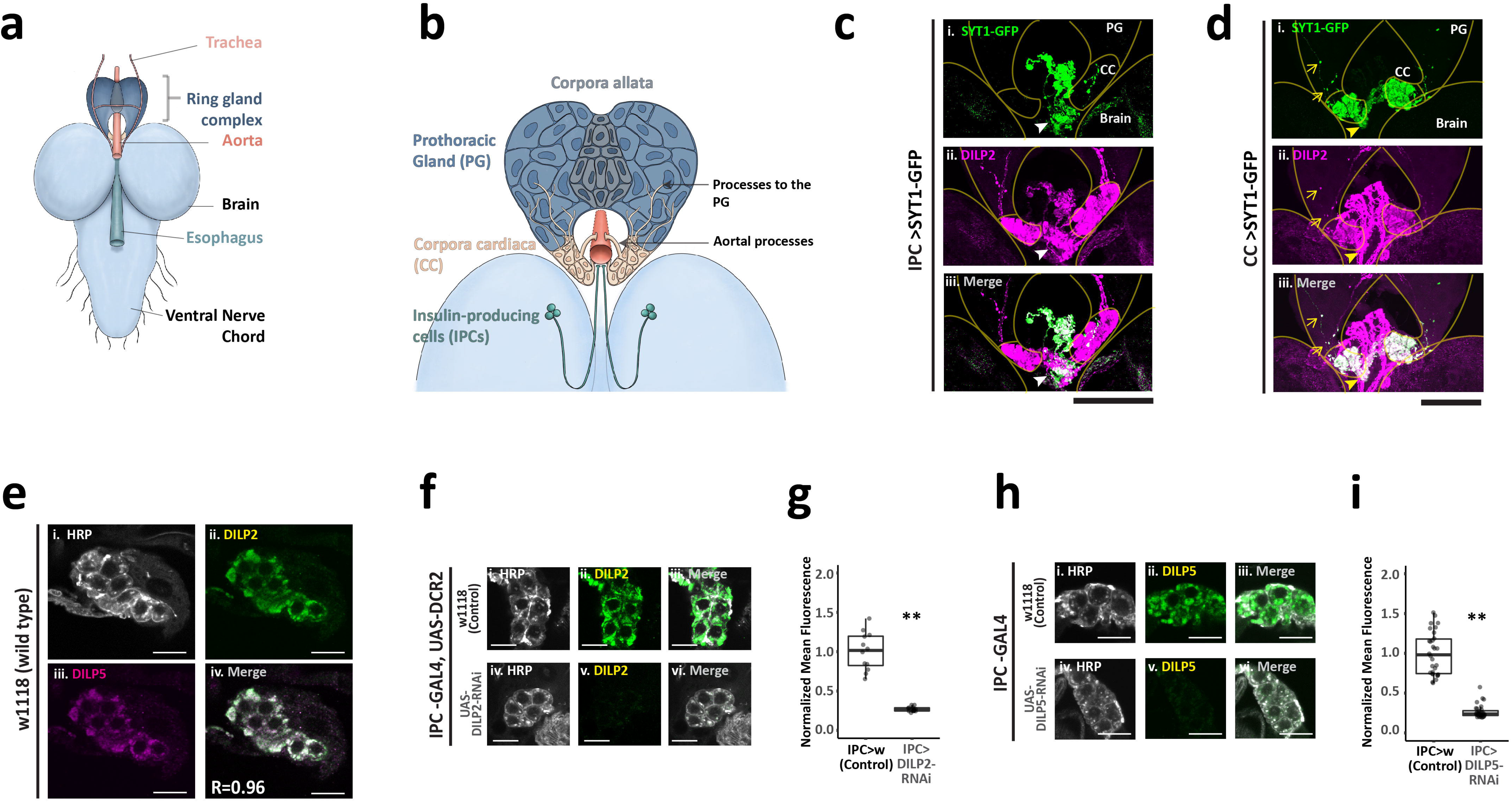
IPC-derived DILPs 2 and 5 accumulate in the corpora cardiaca. (**a-b**) Neuroanatomy of the larval central nervous system and ring gland complex showing (b)innervation of insulin-producing cells(IPC) and the corpora cardiaca(CC). (**c-d**) Maximum z-volume projection of brain and ring gland complex of larvae expressing Synaptotagmin-1-GFP (**c.i and d.i, green**) specifically in the (c) IPC (*dilp2-*gal4) or the (d) CC (*akh-*gal4). Tissues are co-stained for DILP2 (**c.ii and d.ii, magenta**). **Solid arrows** denote aortal projections from (c) IPC (white arrow) and (d) CC (**yellow arrow**). **Open arrows** (yellow) show CC projections to the PG. Merged channels (c.iii and d.iii) channel show colocalization. Scale bar = 50um (**e**) Wild-type CC soma stained for neuronal marker HRP (**e.i; gray**), DILP2 (**e.ii, green**) and DILP5 (**e.iii, magenta**). Merged channel (**e.iv**) show DILP2 and DILP5 colocalization. R = pixel-based Pearson’s correlation coefficient between the same. Scale bar = 10um. (**f,h**) CC soma of IPC-specific(*dilp2*-gal4) (f)DILP2 RNAi in Dicer2-expressing background or (h)DILP5 RNAi (**h.iv-vi**) together with wild-type controls(**f.i-iii and h.i-iii**). Stained for neuronal marker HRP(**f.i, f.iv, h.i and h.iv, gray**) and (**f**)DILP2 (**f.ii and f.v, green**) or (**h**)DILP5 (**h.ii and h.v, green**). Merged channels (f.iii, f.vi, h.iii and h.iv) for clarity. Scale bar = 10um. (**g, i**) Average fluorescence intensity of (**g**)DILP2 or (**i**)DILP5 normalized to the control mean. Each point represents a single CC soma. n =3-7; **= p<0.01, Student’s t-test.

In order to clarify the presence and production of Dilp2 in CC cells, we co-labelled the IPC projections by over-expressing a GFP-tagged Synaptotagmin-1 (*SYT1-GFP*) using a *dilp2-gal4* driver (*IPC>SYT1-GFP*, Figure 1c) and anti-Dilp2 immunostaining. We confirmed the presence of IPC processes containing DILP2 and innervating the aorta (Figure 1c). Additionally, Dilp2 could also be detected without GFP colocalization in the CC soma (Figure 1c). Expressing *SYT1-GFP* using a CC-specific GAL4 driver (*akh-gal4*), we confirmed that Dilp2 indeed co-localizes with GFP in CC cells and their processes projecting on the PG and the aorta (Figure 1d). CC soma also stain for anti- Dilp5, and both Dilp2 and Dilp5 show high degree of subcellular colocalization (Figure 1e), suggesting that these two Dilps may serve a common function in the CCs. Their presence in the soma of the CC cells could either be due to a transient expression in the CCs or their uptake by the CCs from a pool initially produced by the IPCs. To address this point, we specifically knock-down *DILP2* or *DILP5* expression in the IPCs. This led to their loss in CC cells (Figures 1f-i), indicating that the IPCs are the only source of Dilp production, followed by their specific uptake in the CCs. Of note, the knock-down of Dilp5 expression in the IPCs do not affect the uptake of Dilp2 by the CCs (Figure S1).

### IMPL2 mediates the uptake of Dilp2 and Dilp5 in the Corpora cardiaca

In addition to the canonical insulin receptor (dInR), *Drosophila* larvae express another insulin-binding protein called IMPL2. IMPL2 binds circulating Dilp2 and 5, resulting in the downregulation of insulin/IGF-signalling (IIS) in target tissues (Figueroa-Clarevega & Bilder, 2015). Beside tumorous imaginal discs and fat body (Figueroa-Clarevega & Bilder, 2015; Lee et al., 2018), which produce a circulating form of ImpL2, a subset of neurons in the larval brain express the IMPL2 gene (Honegger et al., 2008). Using specific anti-ImpL2 antibodies, we also detected IMPL2 in the CCs (Figure 2a). Similar staining patterns between IMPL2 and Dilp2 suggests that they may be partially bound inside the CC. Staining for IMPL2 in Dilp2/Dilp5 double knock-out larvae ruled out a possible staining artefact due to cross-reactivity of the respective antibodies (Figure S2). Additionally, our results show that the presence of IMPL2 in the CC is independent of Dilp2 and 5 production (Figure S2).

**Figure 2.**
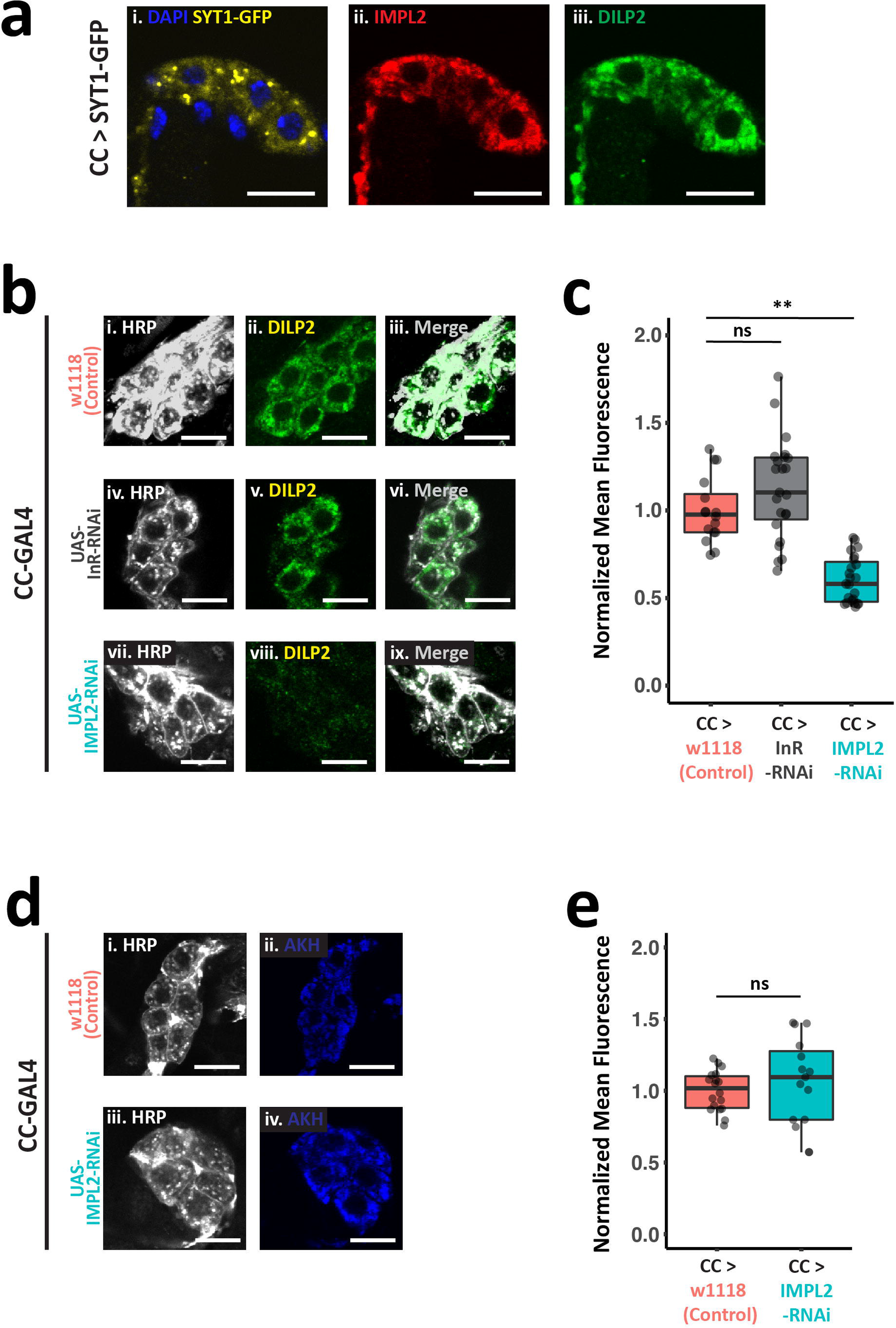
IMPL2 in the corpora cardiaca is necessary for DILP uptake. (**a**) CC soma specifically expressing(*akh-*gal4) Synaptotagmin-1-GFP (**a.i, yellow**); stained for nuclei (**a.i,blue**), IMPL2 (**a.ii, red**) and DILP2 (**a.iii, green**). (**b**) CC-specific(*akh-*gal4) RNAi-mediated knockdown of insulin receptor (**b.iv-vi**), IMPL2 (**b.vii-ix**) and wild-type control (**b.i-iii**). Stained for neuronal marker HRP (**b.i, b.iv, and b.vii, gray**) and DILP2 (**b.ii, b.v and b.viii, green**). Merged channels (**b.iii, b.iv and b.ix**) for clarity. (**d**) CC-specific(*akh-*gal4) RNAi-mediated knockdown of IMPL2 (**d.vii-ix**) and wild-type control (**b.i-iii**). Stained for neuronal marker HRP (**b.i, b.iv, and b.vii, gray**) and AKH (**d.ii,and d.iv, blue**). (**c, e**) Average fluorescence intensity of (**c**)DILP2 or (**e**)AKH normalized to the control mean. Each point represents a single CC soma. n =3-5; **= p<0.01, ns = p>0.01; Student’s t-test. Scale bar = 10um.

IMPL2 binds to Dilps but, unlike dInR, does not have a trans-membrane domain (Roed et al., 2018). A previous study (Bader et al., 2013) showed that a membrane-associated form of IMPL2 produced in a few brain neurons binds Dilp2 and facilitates its contact with dInR, resulting in activation of intracellular IIS signalling. While dInR is recycled to the membrane after endocytosis, the IMPL2-Dilp2 complex is then degraded (Bader et al., 2013). In order to clarify the mechanism of Dilp uptake by the CCs, we knocked-down dInR and IMPL2 and compared the amount of Dilps present in CC cells. dInR and IMPL2 knock-down were first controlled for their efficacies in the wing disc and CC respectively (Figure S4 and S5a). While silencing *dInR* in CCs had no effect on Dilp2 localization, *ImpL2* knock-down significant reduced Dilp2 staining in the CC soma (Figures 2b and 2c). Similar results were obtained for Dilp5 localisation (Figure S3). The knock-down of *dInR* reduces CC cell area and shows a slight increase in mean intensities of Dilp2 (Figures 2c) and 5 (Figures S3). Silencing Impl2 in CCs did not affect expression of Impl2 in brain neurons (Figures S5b). We conclude from these experiments that CC cells require IMPL2, but not dInR, to take up Dilps. We next ensured that preventing Dilp uptake in the CC does not affect endogenous glucagon-like AKH production by staining CCs with anti-AKH antibodies (Figures 2d and 2e). Therefore, Impl2 silencing in CC cells is an efficient tool to study the physiological function of the pool of Dilps present in the CCs.

### Corpora cardiaca act as a functional Dilp store

Larvae lacking Dilp2 and Dilp5 show reduced growth during larval stages, require extended time to develop and produce smaller adults (Grönke, Clarke, Broughton, Andrews, & Partridge, 2010). To evaluate the contribution of the CC store of Dilp2 and Dilp5 to general IIS, we measured the developmental time, larval growth and adult size of animals devoid of Impl2 in CC cells, therefore lacking Dilp accumulation in CCs. We observed that larvae of the *CC>Impl2-RNAi* genotype show approx. 10 hr advanced pupariation compared to controls (Figure 3a). However, pupal volume, as a measure of total larval growth, adult mass and wing area are not modified (Figure 3b-d). This indicates that animals devoid of a CC-Dilp store experience accelerated growth without changing final size. Measuring larval mass across the L3 stage, we indeed found that removal of CC-Dilps leads to an increase in larval growth rate (Figure 3e). Accelerated growth was previously observed upon general increase in intracellular IIS (Brogiolo et al., 2001) or over-expression of *DILP2* (Géminard, Rulifson, & Léopold, 2009; Ikeya et al., 2002) during larval development. When measuring circulating Dilp2 from *CC>Impl2-RNAi* larvae, we found it doubling compared to controls (Figure 3f). To support this finding, we also found elevated levels of fat body IIS at 104hr AEL compared to control genotypes, measured as the relative distribution of the transcription factor dFOXO between nucleus and cytoplasm (Figure 3g and 3h). Therefore, preventing DILP uptake in the CC increases circulating Dilp2 levels, resulting in accelerated growth.

**Figure 3.**
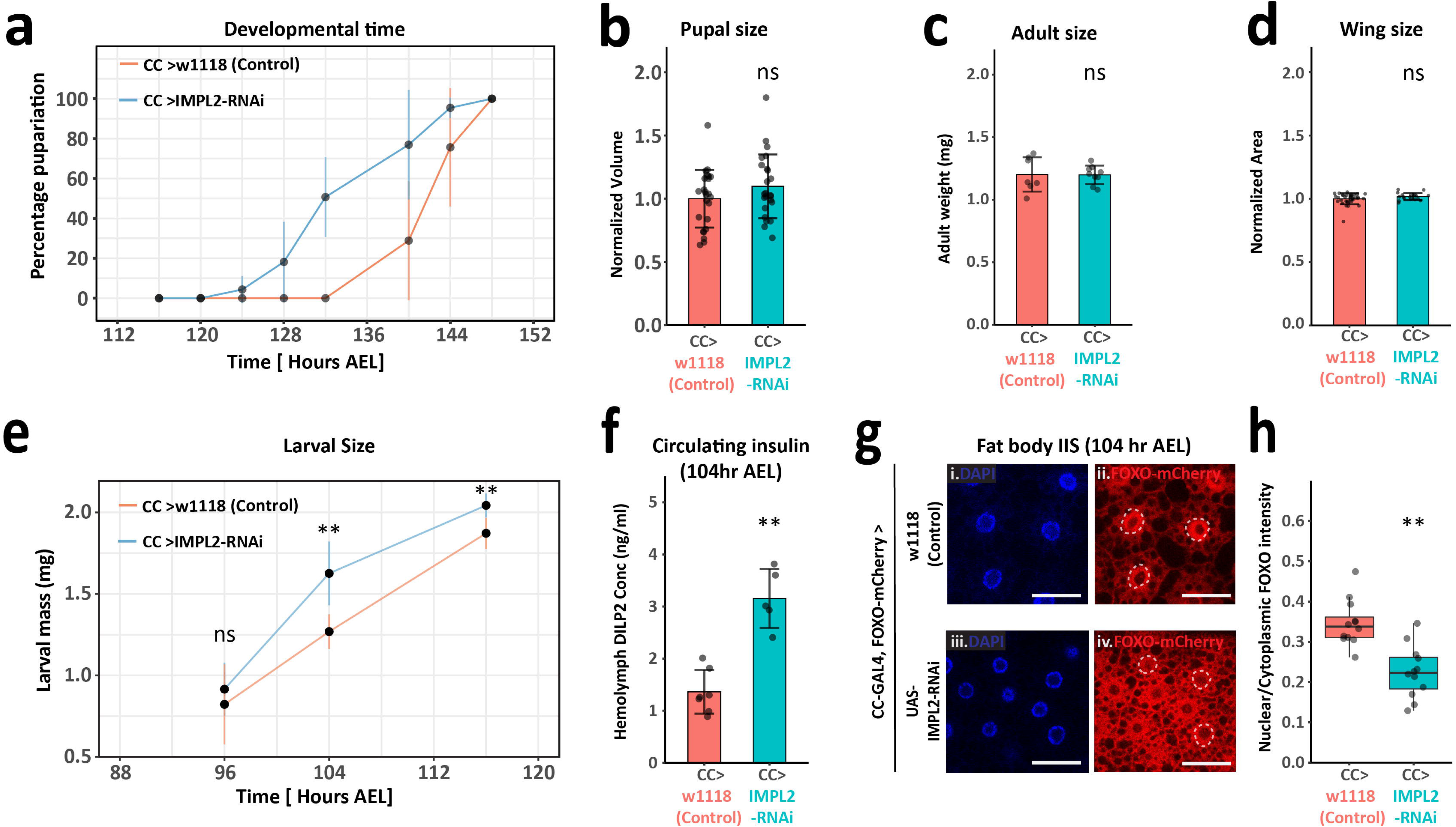
Corpora cardiaca sequesters circulating DILPs. (**a**) Percentage pupariated of indicated genotype at various timepoints. AEL= after egg-laying. n=6 for each genotype. (**b**) Pupal volume (n= 24), (**c**) adult weight (n=7, 9) and (**d**) adult wing area (n=55, 20) normalized to control. Each point represents a larva/fly in the case of pupal volume or wing area, and mean of a cohort of 8-15 flies in the case weight measurements. (**e**) Larval mass of indicated genotype at specified developmental time (n=8-12). Each point represents mean of cohorts of 13-15 flies. (**f**) DILP2-HA-FLAG concentration in larval hemolymph of indicated genotype using driver line *akh*-gal4,dilp2-HF (n=7,5). (**g**) Fat body of indicated genotype stained for nuclei (**g.i and g.iii, blue**) and mCherry (**g.ii and g.iv, red**). White dashed outlines show the position of some nuclei in the mCherry channel. Scale bar = 50um. (h) Boxplots showing ratio of nuclear to cytoplasmic mCherry intensity in fat bodies. Lower ratio indicates higher insulin activity. (n= 10) Bar plots show mean and error bars indicate standard deviation. AEL= after egg-laying. **= p<0.01, ns = p>0.01, (**b-e**) Student’s t-test and (**f,h**) two-tailed Mann-Whitney U test.

### CC-Dilps are packaged for co-secretion with AKH

Our previous results indicate that approximately equal amounts of Dilp circulate in the hemolymph and are sequestered in the CC cells (Figure 3f). We therefore studied the control of secretion of this major pool of CC-Dilp. Based on colocalization using light microscopy, it was previously proposed that in ImpL2-expressing neurons, ImpL2/Dilp2 complexes are degraded through the late endosomal route (Bader et al., 2013). Given the small size of the CC soma, we used electron microscopy to identify DILP-containing structures inside the CC cells (Figure 4a). Stitching several high-resolution images from a single tissue section shows the overall organization of the CC cytoplasm (Figure 4b). The cytoplasm contains large networks of ER, occasionally interspersed with Golgi complexes, and clusters of dense-core secretory vesicles (DCVs) occupying the perinuclear space (Figure 4b). This architecture is in line with the capacity of CC cells to secrete the glucagon-like hormone AKH, and similar observations have been made in other insects (Willey & Chapman, 1960). When probed for Dilp5 using immuno-gold staining, CC cells showed high reactivity in clusters of DCVs compared to rest of the cytoplasm (Figures 4c and 4d). Probing simultaneously for Dilp2 and 5 using gold particles of two different sizes, we observed a co-localization of both Dilps in DCVs, but also in multi-vesicular bodies (MVBs) (Figure 4e). MVBs form in the late endocytic pathway and are responsible for recycling and degradation of endocytic compartments (Cosker & Segal, 2014; Sorkin & Von Zastrow, 2009). Alternatively, they also give rise to secretory granules (Gondré-Lewis, Park, & Loh, 2012). Our results therefore suggest that Dilp2 and 5 are taken up and packaged into secretory DCVs, making the CC cells a potential alternative source of insulin in the *Drosophila* larva.

**Figure 4.**
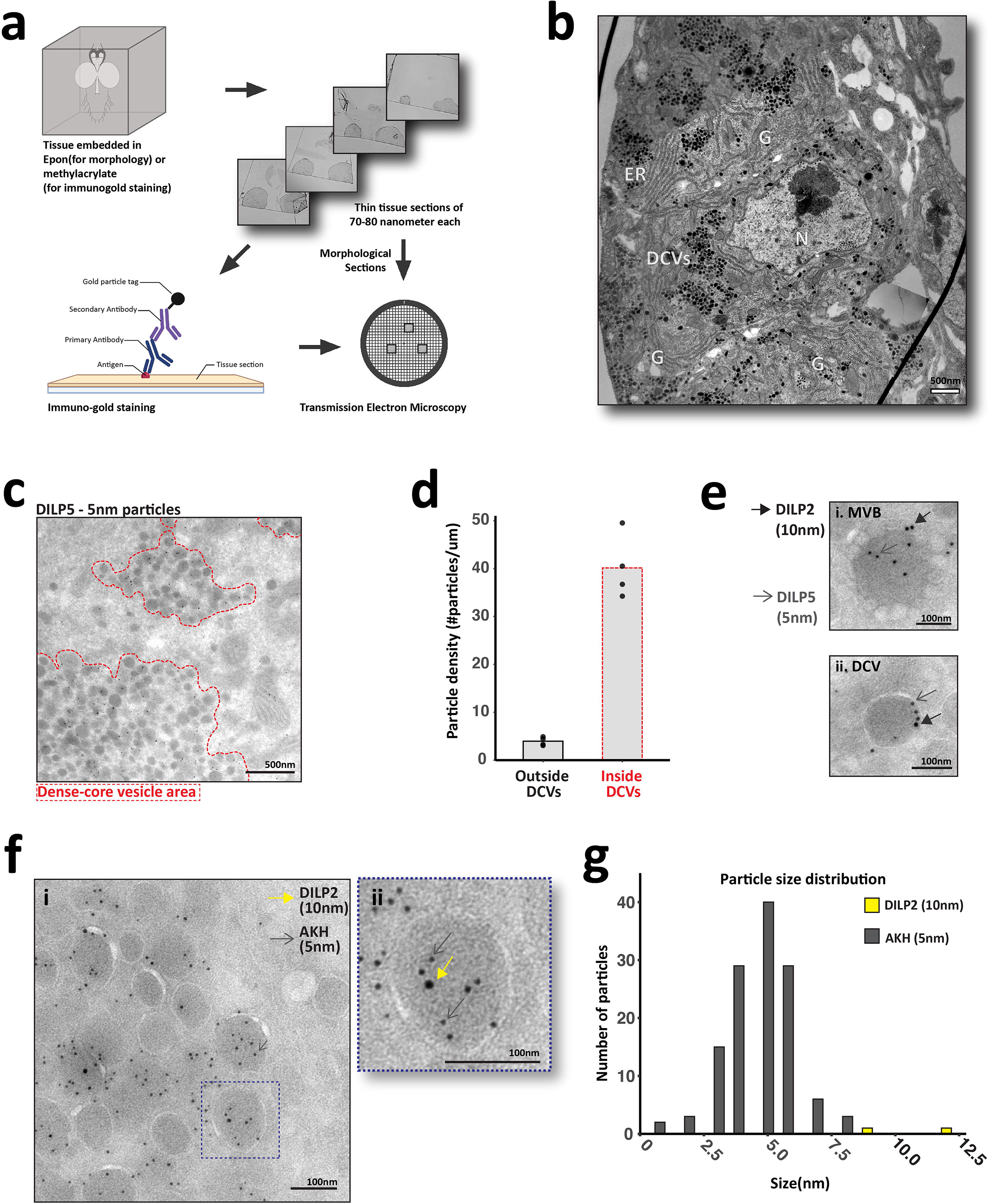
DILPs and AKH colocalize in corpora cardiaca dense-core vesicles. (**a**) Schematic representation of experimental approach. Electron micrograph of (**b**) the CC soma (epon-embedded sections). N = nucleus, G = Golgi complex, ER = endoplasmic reticulum, and DCV= dense-core vesicle. Scale bar = 500nm. (**c**) Part of cytoplasm in the CC soma (methylacrylate sections). DILP5 labelled with immunogold particles of 5nm diameter. Dashed red line indicates area occupied by DCVs. Scale bar = 500nm (**d**) Quantification of gold particle density (no. of particles/um^2^) from DCV area (**dashed red line**) or rest of the cytoplasm (**solid black line**) from multiple acquisitions (n=4). Bar plot indicates mean. High-resolution electron micrographs of (**e**) a multi-vesicular body (**MVB, e.i**) and a dense-core vesicle(**DCV, e.ii**) in the CC soma. DILP2 and DILP5 are stained by 10nm (**closed arrow**) and 5nm (**open arrow**) immunogold particles respectively. Scale = 100nm. (**e**) DCV area (**f.i**) stained for DILP2 (**10nm gold particle**) and AKH (**5nm gold particle**). Dashed blue box outlines a single DCV (**f.ii**) magnified on the right, DILP2 (**closed yellow arrow**) and AKH (**open gray arrow**). (**g**) Scale bar = 100nm. Histogram showing immunogold particle size vs. frequency distribution in (**f**), highlighting relative abundance of DILP2 (**yellow**) and AKH (**gray**).

CC cells endogenously produce the glucagon-like peptide hormone AKH and release it under nutrient-limiting conditions (S. K. Kim & Rulifson, 2004). When probing CC cells for AKH and Dilp2 using two differently-sized gold particles, we found that AKH and Dilp2 co-localize in the same DCVs (Figure 4f). We also noted that AKH seems more abundant than Dilp2 in these DCVs (Figure 4g). Therefore, two functionally antagonistic hormones, AKH and Insulins, are co-packaged in the same secretory structures, raising the intriguing possibility that they are released by CC cells in the same physiological context (i.e. limiting nutrients).

### CC-Dilps control the time to pupariation under nutrient limitation

What could be the physiological role of the pool of CC-insulin? During the late larval period, animals pass through a nutrient restriction checkpoint (NRC), beyond which larvae are committed to pupal development even when fully starved (Pan, Neufeld, Connor, et al., 2019). Early observations (Beadle et. al., 1938) indicate that when starved after the NRC, *Drosophila* larvae pupariate earlier than fed controls, as we observed for *CC>w1118* (Figure 5a,b).

**Figure 5.**
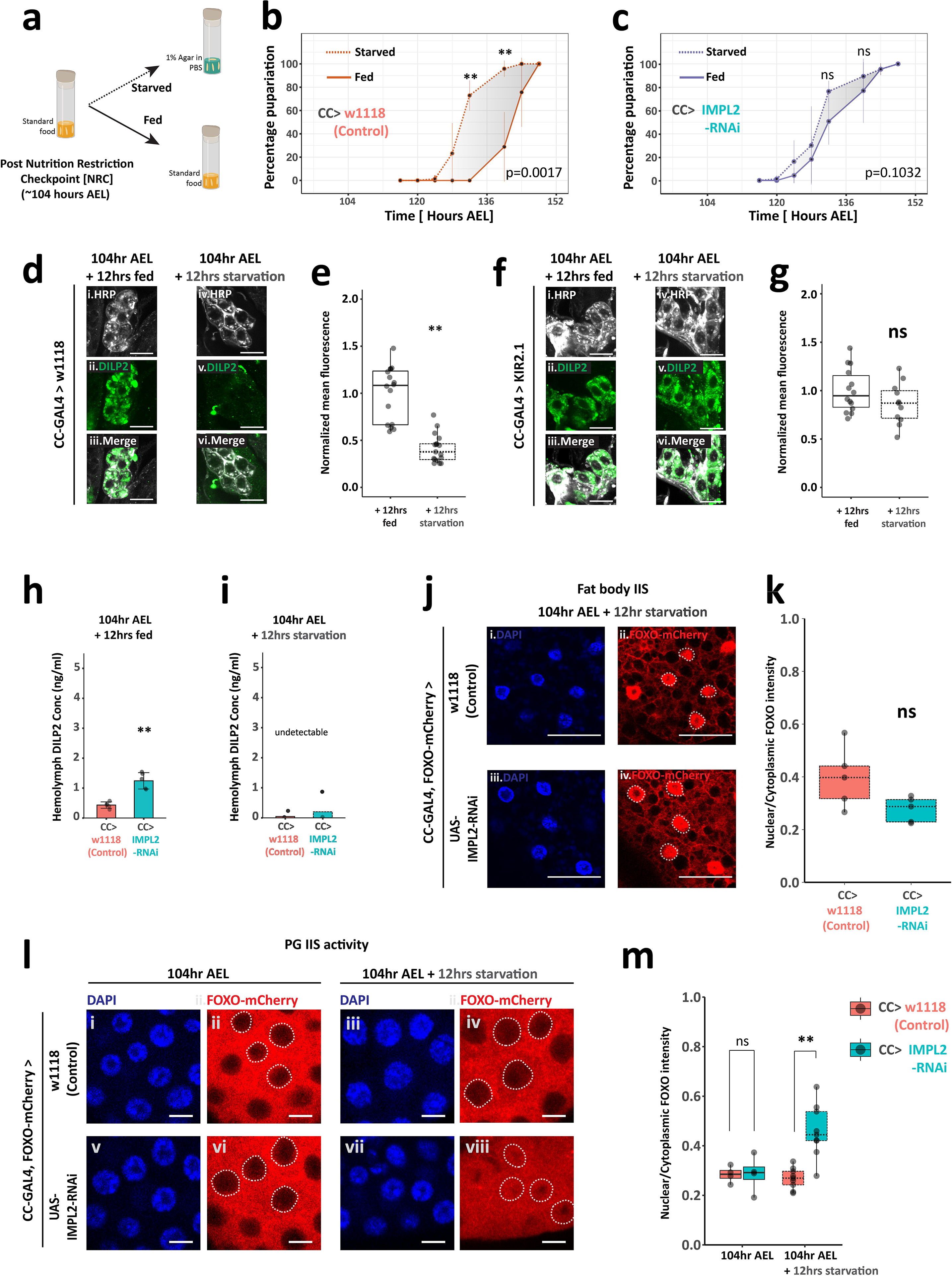
Corpora cardiaca DILPs enable starvation-induced accelerated pupariation. (**a**) Experimental setup for larval starvation (**b-c**) Means of percentage pupariation connected across timepoints for the indicated genotype-starved (**dashed line**) or kept feeding (**solid line**) as shown in (**a**). Fed controls are the same as shown in Figure 3a. n=6 for each curve; p-value from two-way ANOVA with repeated measures and Bonferroni’s post t-test for multiple comparisons. Error bars indicate +/− S.D. **= p<0.01 and ns = p>0.01 (**d,f**) CC soma of genotype (**d**)*akh>*w1118 and (**f**)*akh>*KIR2.1 stained for HRP(**d.i,d.iv, f.i and fiv; gray**) and DILP2 (**d.ii, d.v, f.ii and f.v; green**). KIR2.1 expression inactivates neurons through hyperpolarization. Scale bar = 10um. (**e,g**) Average fluorescence intensity of DILP2 normalized to the mean of fed controls. Each point represents a single CC soma. n =3-5. (**h-i**) DILP2-HA-FLAG concentration in larval hemolymph of indicated genotype using driver line *akh*-gal4,dilp2-HF (n=4-5). (**j**) Fat body of indicated genotype stained for nuclei (**j.i and j.iii; blue**) and mCherry (**j.ii and j.iv; red**). White dashed outlines show the position of some nuclei in the mCherry channel. Scale bar = 50um. (**k**) Boxplots showing ratio of cytoplasmic to nuclear mCherry intensity in fat cells (n=5). (l) Prothoracic gland of indicated genotype stained for nuclei (l.i, l.iii, l.v and l.vii; blue) and mCherry (**l.ii, l.iv, l.vi and l.viii; red**). White dashed outlines show the position of some nuclei in the mCherry channel. Scale bar = 10um. (**k**) Boxplots showing ratio of nuclear to cytoplasmic mCherry intensity in prothoracic gland cells. n=4-8. AEL= after egg-laying. **= p<0.01, ns = p>0.01, (e-g) Student’s t-test and (h,i,k,m) two-tailed Mann-Whitney U test.

Several lines of evidence suggest that the pool of CC-Dilps induces early pupariation upon starvation post-NRC. First, we noticed that, when comparing fed and starved conditions, the acceleration observed for *CC>w1118* controls is absent in the *CC>ImpL2-RNAi* background (Figure 5c). As described before (see Figure 3a,e), fed *CC>ImpL2-RNAi* larvae pupariate earlier than fed *CC>* controls due to accelerated growth, but in *CC>ImpL2-RNAi* larvae, the difference between fed and starved conditions is abrogated (marked as a grey area in Figures 5b,c). We then confirmed that CC-Dilps are released upon low/no nutrient conditions (Figure 5d-g), reminiscent of what is observed for the control of AKH secretion (J. Kim & Neufeld, 2015). The transition into pupal development requires a surge of ecdysone production and previous work indicate that activation of IIS is required for ecdysone biosynthesis by the PG (Boulan, Martín, & Milán, 2013; Colombani, 2005). Therefore, our observation that CC cells send projections to the PG raises the possibility of a local delivery of Dilps to the PG in response to starvation. To test this idea, we compared the levels of circulating Dilp2 in *CC>* and *CC>ImpL2-RNAi* larvae under different nutritional conditions. In fed conditions, suppressing the CC-Dilp pool leads to an increase in circulating Dilp2 (Figure 5h). However, during starvation post-NRC, no circulating Dilp2 is detected either in *CC>* or *CC>Impl2* conditions (Figure 5i), indicating that the release of the CC pool of Dilp2 does not contribute to increasing systemic Dilp2. In parallel, we measured the nucleus/cytoplasm ratio of dFOXO in fat cells as a marker of peripheral IIS (Figure 5j,k). Again, no difference in adipose IIS was observed between *CC>* and *CC>ImpL2-RNAi* larvae starved after the NRC, indicating that under these conditions, CC-Dilp2 does not contribute to general IIS. In contrast, when conducting a similar analysis of the cells of the prothoracic gland, which synthesizes ecdysone, we observe that starvation after the NRC induces an increase in nuclear localization of dFOXO in the PG of *CC>ImpL2-RNAi* but not of *CC>* control animals (Figure 5l,m). This result indicates that CC-Dilps contribute to maintaining a high level of IIS activation in PG cells in response to post-CW starvation. In line with our finding that *CC>ImpL2-RNAi* larvae pupariate earlier under these conditions, we conclude that the pool of CC-Dilps contributes to a local activation of IIS in PG cells, maintaining their capacity to synthesize ecdysone and allowing a premature transition into pupal development in response to late starvation.

## Discussion

In this study, we re-evaluate a previous observation whereby glucagon-producing cells take up insulin peptides, and demonstrate the relevance of this phenomenon during *Drosophila* larval development. In the course of early studies on *Drosophila* insulin-producing cells (S. K. Kim & Rulifson, 2004), it was indeed noted that larval CC cells strongly label for Dilp2 peptides. This puzzling observation suggested that the larval CC could serve the role of a so-called “neurohemal” organ, storing Dilps for delivery in the general circulation under specific conditions. However, the mechanistic and functional aspects of this Dilp store were unknown. Our study now sheds new light on the role of CC-Dilps, linking them to another unexplained observation, the starvation-induced, premature transition of *Drosophila* larvae into pupal development (Beadle et. al., 1938).

### Accelerating development upon post-critical weight starvation

*Drosophila* populations can suffer episodes of nutrient limitation, particularly during the larval period when mobility of individuals is limited, preventing them from exploring alternative food sources. When larvae endure full starvation passed the so-called nutrient restriction checkpoint (NRC), an acceleration of pupariation is observed and animals transit faster to an adult reproductive state, with a trade-off on individual size. This contrasts with the observation that Dilp secretion is inhibited by starvation (Géminard et al., 2009), leading to a reduction of circulating hormone levels. Such reduction should indeed affect the production of ecdysone, which relies on IIS in PG cells (Colombani, 2005). Our present work indicates that upon limiting nutrients, CC cells secrete stored Dilps to the PG in a paracrine fashion (through PG-specific projections) without affecting systemic IIS. The CC-derived Dilps sustain IIS in PG cells, providing the necessary signalling input for ecdysone production and accelerated pupariation. Indeed, IIS activation could prevent starvation-induced autophagy in PG cells, which has been shown to block ecdysone production in condition of a pre-NRC starvation, by shunting cholesterol away from the biosynthetic pathway (Pan, Neufeld, & O’Connor, 2019). Recent evidence suggests that alternative nutrient-independent growth factors such as Jelly-belly (Jeb) can also act on PG cells to stimulate IIS and ecdysone production (Pan & O’Connor, 2020). Jeb and Dilps could constitute separate inputs on ecdysone production, explaining why removal of CC Dilps only affects acceleration upon nutrient restriction but does not delay pupariation. Whether CC-Dilps are uniquely targeted to the PG, and through which mechanism, are still open questions.

### Storing Dilps away from general circulation

We find that preventing CC-Dilp storage increases circulating levels by 2-fold. This accelerates animal’s growth rate during the late larval stages, when circulating Dilp levels wane (compare Figures 3f and 5h; also see Okamoto & Nishimura, 2015). We also noted that such accelerated larval growth does not affect adult size due a compensatory advance in pupariation. This result suggests an interdependency relationship between the two Dilp reservoirs, whereby the CC-Dilp store forms from ILP-produced insulins that are derouted from the general circulation in conditions of optimal nutrition. It has been described previously that the IPC project their termini in proximity of the CCs onto the aorta. Whether CC-Dilp accumulates through local delivery of IPC-produced Dilps or by uptake from the general circulation is still unknown. Insulins bind to their cognate receptor InR in target tissues and are then co-endocytosed and later degraded (Sopko & Perrimon, 2013; Sorkin & Von Zastrow, 2009). Our study demonstrates that the uptake of Dilps by CC cells does not rely on InR but rather Imp-L2, an alternative insulin-binding protein previously shown to bind circulating Dilps. This result contrasts with the previously described case of Imp-L2-producing *Hugin* neurons in the larval CNS, where both InR and Imp-L2 are required for Dilp uptake (Bader et al., 2013). This difference might have important functional consequences. In contrast to endocytosed Dilps/Inr complexes, which are prone to degradation, endocytosed ImpL2/Dilp complexes are routed towards storage in DCV, in the absence of local activation of IIS. Dilps are co-packaged with AKH in DCVs, possibly during the DCV maturation phase. This phase is characterized by cargo exchange between endo-lysosomal compartments and immature vesicles budding off from the trans-Golgi network (Topalidou et al., 2016). Recent studies (Gondré-Lewis et al., 2012; Lund, Lycas, Schack, Andersen, & Gether, 2020) suggest that RAB2 and associated proteins are involved in this exchange. However, we do not know if these components participate in routing DILPs and AKH to DCVs in the CC. Further, Dilps are anterogradely transported along the axonal projections in the PG. This process is similar to the transcytosis of various trophic factors and pathogens observed in mammalian nervous systems (Von Bartheld, 2004). Unlike simultaneous uptake and release previously demonstrated in other neuron types (Hémar, Olivo, Williamson, Saffrich, & Dotti, 1997; Von Bartheld, Wang, & Butowt, 2001; Yamashita, Joshi, Zhang, Zhang, & Kuruvilla, 2017), we show here that Dilp uptake and release are temporally separated during larval development. In all, this sheds light on the unique ability of CC neurosecretory cells to convert an endocrine signal like insulin into a paracrine one.

In conclusion, we reveal here a mechanism of organ sparing relying on the storage and release of insulins, allowing the maintenance of IIS activation and ecdysone production in a paracrine organ. The existence of such mechanism opens the intriguing possibility that local pools of insulin/IGF could be used in response to environmental signals to maintain IIS signalling in spared organs in other models.

## Acknowledgements

This article is dedicated to the memory of our great colleague and mentor Prof./Dr. Suzanne Eaton. We are grateful to Dr. Natalie Dye for her supervision and leadership. We thank Dr.Jonathan Rodenfels, the Eaton and Leopold lab members for insightful discussions and comments on the manuscript. We would also like to thank the BDSC, Prof. Linda Partridge and Dr. Jason Tennessen for providing us with fly lines, Prof. Marc Tatar and Dr. Stephanie Post for sharing reagents, and Dr. Hugo Stocker for gifting us the Imp-L2 antibody. The scientific illustrations were done by Dr. Bertsy Goic. This work was supported by Deutsche Forschungsgemeinschaft (DFG/FOR2682), Max-Planck-Gesellschaft and Institut Curie, INSERM.

## Author contributions

Conceptualization, S.G., S.E., P.L., Methodology, S.G., S.E., P.L., Investigation, S.G., W.L., M.W.-B., Writing and editing, S.G., P.L., Funding Acquisition, S.E.; Supervision, S.E., P.L.

## Materials and methods

### Flies

All fly lines/crosses were maintained and grown on standard media containing cornmeal, molasses, agar and yeast extract under a 12hour light/dark cycle at 25°C (unless explicitly stated otherwise).

List of fly strains used.

List of fly strains used
| Strain | Source | Identifier |
| --- | --- | --- |
| w1118 | BDSC | #5905 |
| IPC-Gal4 (Dilp2-Gal4) | BDSC | #37516 |
| CC-Gal4 (akh-Gal4) | (Tennessen, Baker, Lam, Evans, & Thummel, 2011) | NA |
| dFOXO-mCherry | (Kakanj et al., 2016) | NA |
| gDilp2-HA-FLAG | (Park et al., 2014) | NA |
| UAS-SYT1-GFP | BDSC | #6926 |
| UAS-KIR2.1-GFP | BDSC | #6596 |
| UAS-Dcr2 | BDSC | #24651 |
| UAS-DILP2-RNAi | BDSC | #31068 |
| UAS-DILP5-RNAi | BDSC | #31378 |
| UAS-InR-RNAi | BDSC | #31037 |
| UAS-IMPL2RNAi | BDSC | #64936 |
| DILP-2,3,5,7 knock-out | BDSC | #30889 |
| ap-Gal4, tub-Gal80TS | Spannl et. al., 2020 | NA |

Fly strains *dilp2-Gal4,>dcr2* , *akh-Gal4,Foxo-mCherry* , *akh-Gal4,qdilp2-HF* and *ap-Gal4,tub-Gal80TS,Foxo-mCherry* were generated in this paper by crossing the above lines and are available upon request.

### Developmental timing and starvation

Fly crosses were fed fresh yeast paste and allowed to lay eggs for 3 hours on apple-juice agar plates. Following egg-laying, the plates were stored at 25°C. After 18-24 hours of incubation, freshly hatched L1 larvae were collected in PBS in intervals of 2 hours. In order to prevent overcrowding, no more than 20-30 larvae were transferred into a single food vial using a micropipette and shifted to the incubator at 25°C with 12/12 day-night cycle. At specified time after egg-laying, larvae were carefully washed out of the food using PBS. Subsequently they were quickly transferred to 1% agar vials (same quantity as food). Fed controls were not washed out, and left to feed in their original vials. Both starved and fed control vials were returned to the same incubator.

For pupariation timing, number of pupae including white pre-pupae were manually counted at fixed intervals as indicated. Counting was stopped when there was no increase for 3-4 consecutive time points. The same dataset have been used in Figure 3a, and to depict fed controls in Figure 5b,c. For all other starvation experiments, starved and fed control larvae were washed out of the agar/food with PBS, rinsed and used in respective assays.

### Weight measurements

#### Larval weight

At specified developmental stage, larvae were collected from food/agar in PBS, washed 3x times to remove stuck food and cold-anesthetized. Female larvae were selected under the microscope to prevent sex-dependent size variation. Cohorts of 13-15 larvae were blotted on a tissue paper (to ensure no extra liquid) and transferred to a pre-weighed 1.5ml micro-centrifuge tube using forceps. The tube with larvae was then weighed using a fine balance (Sartorius).

#### Adult weight

Virgin females (2 days after eclosion) were anesthetized and collected in pre-weighed 1.5ml micro-centrifuge in cohorts of 10 flies per tube. The tube with adult flies was weighed using a fine balance (Sartorius).

Average weight in each case was calculated by dividing the weight difference with the number of larvae/flies in each tube.

### Size determination

#### Adult wing area

Females (1 day after eclosion) were collected and stored in isopropanol. Prior to dissection, flies were transferred to a cavity dissection-glass. Care was taken to keep flies submerged in adequate isopropanol at all times. One wing from each fly was dissected using fine forceps (Dumont, No.5-SA) and transferred to a glass slide along with a small amount of isopropanol (in forceps due to capillary action). It is important to keep wings flat on the slide surface. This can be achieved by repeatedly moistening the wings with isopropanol, using forceps. Once the desired number of wings were obtained, 30ul of Euparal (Roth, cat#7356.1) was pipetted on the slide. Next, a glass coverslip (22mm/22mm/17um) was placed covering the entire area on the slide occupied by wings. To ensure absence of air bubbles and minimum displacement in the wings, it is important to place the coverslip starting from one edge. Following mounting, the slides were left to dry for 2 hours prior to imaging. Single wing images were taken using Zeiss Sv11 dissection scope fitted with a CCD camera (Jenoptik ProgRes C10).

#### Pupal volume

Pupae (24-30-hours APF) were collected using a wet brush. Following which, they were washed with PBS to dislodge attached food and glue. Each pupa was then transferred to a single well of a transparent 24-well plate and imaged using Zeiss Sv11 dissection scope fitted with a CCD camera (Jenoptik ProgRes C10). Sex determination was done post-eclosion and only females were taken for analysis.

### Dilp2-HF ELISA

Nunc MaxiSorp Flat-Bottom plates (Thermo Fisher Scientific) were coated with anti-FLAG M2 antibody (Sigma, cat#F1804) diluted in BupH Carbonate-Bicarbonate Buffer (Thermo Fisher Scientific, cat#28382) overnight at 4°C. The plates were washed 2x in wash buffer – PBS + 0.2% Tween20 (Sigma, cat#P7949) and blocked in 2% filter-sterilized BSA in PBS overnight at 4°C. 1mL of hemolymph was collected by bleeding 5-8 heterozygous gDilp2-HF larvae on a clean cavity slide. The hemolymph was diluted in 55mL PBS and centrifuged at 10,000rpm for 10 minutes to remove hemocytes and debris. 50mL supernatant and 1mL of anti-HA-HRP antibody (Roche, cat#12013819001) solution (1:500) was incubated in the coated wells overnight at 4°C. 100 ml of 1-Step-Ultra TMB-ELISA Substrate (Thermo Fisher, cat#34028) was pipetted to each well and incubated for 30 minutes in dark at room temperature. Reaction was stopped by adding 100 ml of 2M sulfuric acid per well. The samples colorized instantly and the absorbance was measured immediately at 450nm using a plate reader (BMG Labtech). Standard curve was generated by self-generated FLAG(GS)HA peptide. The detailed method is described in Park et al., (2014).

### Immunofluorescence

Freshly dissected tissue in PBS was fixed with 4% paraformaldehyde (cat#16005, Aldrich) for 20 minutes. After fixation, tissues were thoroughly washed 3-4 times with PBS. The following protocol was used depending on tissue type.

#### Brain, ring gland and fat body

Tissues were permeabilized with PBS containing 0.2%(v/v) Triton X-100(Serva Electrophoresis GmbH, cat#37240.01) for 2 hours at room temperature with shaking. Primary incubation with antibody was done overnight in incubation buffer- PBS with 0.2% Triton X-100 (PBST) and 1mg/ml bovine serum albumin (BSA) (Sigma, cat#A2153). Incubation was followed by 3×15 minute washes with the same incubation buffer without antibodies. Secondary incubation was done with antibody diluted in incubation buffer supplemented with 4%(v/v) normal goat serum (NGS) overnight at 4°C. Tissues were then washed 4X15 minutes in 0.2%PBST and mounted on glass slides using VectaShield H-1000(cat# H-1000-10) mounting medium.

#### Wing discs

Discs were permeabilized with PBS containing 0.05%(v/v) Triton X-100 supplemented with 1mg/ml bovine serum albumin (BSA) and 250mM NaCl for 3X10 minutes at room temperature with shaking. Primary incubation with antibody was done overnight in incubation buffer- PBS with 0.05% Triton X-100 (PBST) and 1mg/ml bovine serum albumin (BSA). Incubation was followed by 3×15 minute washes with the same incubation buffer without antibodies. Secondary incubation was done with antibody diluted in incubation buffer supplemented with 4%(v/v) normal goat serum (NGS) for 2-3 hours. Wing discs were then washed 3X15 minutes in 0.1%PBST and mounted on glass slides using VectaShield H-1000 mounting medium.

All washing and incubation steps were performed on a shaker. Fat bodies from late larval stages disintegrate easily and requires extra care while handling, preferably shaking at a low rpm.

List of primary antibodies.

List of primary antibodies
| Antibody | Host species | Dilution | Source |
| --- | --- | --- | --- |
| Dilp2 | Rat | 1:400 | (Géminard et al., 2009) |
| Dilp5 | Rat | 1:1000 | This paper |
| Dilp5 | Rabbit | 1:500 | (Géminard et al., 2009) |
| Akh | Rabbit | 1:800 | (Isabel,Martin,Chidami, Veenstra,&Rosay, 2005) |
| mCherry | Rabbit | 1:500 | Invitrogen #PA5-34974 |
| GFP | Rabbit | 1:500 | Invitrogen #A111022 |
| IMPL2 | Rabbit | 1:1000 | (Bader et al., 2013) |

Secondary antibodies/dyes used were DAPI (1:2000 from 5mg/ml stock, Roche), DyLight 405 goat anti-HRP (1:500, Jackson IR, cat#123-475-021), AlexaFlour-488 goat anti-rabbit (1:1000, Invitrogen #A11034), AlexaFlour-568 goat anti-rabbit (1:500, Invitrogen #A11036) and AlexaFlour-647 goat anti-rat (1:500, Invitrogen #A21247).

### Confocal Light microscopy

Three imaging setups have been used for fixed samples-

1. Olympus FV1000 laser scanning confocal microscope on Olympus BX61 inverted stand and driven by FV10-ASW 1.7 software. Brain and ring gland whole mounts were imaged with Olympus UApochromat 40x 1.3NA oil immersion objective. Corpora cardiaca cells were imaged with Olympus UApochromat 60x 1.35 NA oil immersion objective.
2. Andor spinning disc confocal microscope on Olympus IX81 inverted stand using Andor iXON 897 EMCCD camera with Yokogawa CSUX1 scanhead, running on Andor software. Some fat bodies were imaged with Olympus U Plan SApo 60x/ 1.35NA oil immersion objective.
3. Olympus FV3000 laser scanning confocal microscope on Olympus IX83 inverted stand and driven by FV31S-SW acquisition software. Corpora cardiaca cells and wing discs were imaged with Olympus UPLXApochromat 60x 1.42 NA oil immersion objective.

### Electron microscopy

#### Morphology

Dissected brain-ring gland complexes were initially fixed in PBS with 2%PFA (cat#16005, Aldrich) and 1% glutaraldehyde (cat#G5882, Sigma) for 2 hours at room temperature followed by overnight incubation at 4°C. Then the samples were post-fixed with 1% OsO4 (cat# 19190, EMS) and 1.5% potassium ferrocyanide (cat#: P9387, Sigma-Aldrich) in water on ice for 1 hour. En bloc staining was done in the dark with 1% UA (cat# 21447, Polyscience Europe GmbH) in water at 4°C overnight. Following infiltration and embedding, 70nm sections were cut on a Leica Ultracut UCT (Leica Microscopy systems) and picked up with formvar (cat# 15800, EMS) coated slot grids (cat# G2010-Cu, EMS). All samples were then post-stained with 1% UA (cat# 21447, Polysciences Europe GmbH) in water for 10 minutes and 0.04% lead citrate (cat# 17800, EMS) for 5 minutes. Electron micrographs were obtained at 80kV using Morgagni (Emsis Morada CCD camera, formerly Sys/Olympus) and 100kV using Tecnai 12 (TVIPS F416 camera) electron microscopes (both from Thermo Fisher Scientific (formerly FEI/Philips)).

#### Immunogold labelling

Dissected brain-ring gland complexes were initially fixed in PBS with 4% PFA and 0.05% glutaraldehyde for 20 minutes at room temperature. Post-fixation was done in 1:1 dilution of the initial fixative overnight at 4°C. Ring glands were then embedded with 12% gelatin (cat# G-2500, Sigma-Aldrich) in PBS and infiltrated with 2.3M sucrose (cat# 21600, EMS) in PBS overnight at 4°C. The sucrose infiltrated Ring gland in gelatin was mounted on an aluminum pin and snap frozen in LN2 and cryosectioned on a Leica Ultracut EM UC6 ultramicrotome (Leica Microscopy systems). 70nm Sections were picked up using a 1 :1 mixture of 2.3M sucrose and 2% methyl cellulose (cat# M-6385, Sigma-Aldrich) as a pick-up solution and sections were transferred to carbon and formvar coated mesh grids (cat# H100-Cu, EMS). After removal of gelatin by incubation on PBS at 37 °C, sections were labelled with rabbit-anti-Dilp2/Dilp5 or goat-anti-AKH Antibody (diluted 1 to 20 or 1 to 100 from original antibody solution) followed by secondary antibody coupled to 10-nm gold (British Biocell or Bbisolutions) and secondary fluorescence antibodies (donkey-anti-rat Alexa594 cat# A-21209, Molecular Probes Europe B.V., goat-anti-rabbit Alexa488 cat# A-21206, Mol. Probes Europe B.V.) The sections were first imaged with epifluorescence on a Zeiss upright ApoTome microscope with Axiocam506 camera to locate regions of interest in individual sections and contrasted with a mixture of 1.9% methyl cellulose/0.3% UA for 10min on ice and the corresponding regions of interest were observed by using a Morgagni or Tecnai12 electron microscope.

### Image analysis and quantification

#### Whole cell

Individual cells from multiple samples were manually segmented and mean intensity was measured using Fiji/ImageJ (Schindelin et al., 2012).

#### Co-localization

Single confocal images from two different channels were used in Coloc 2 plugin in Fiji/ImageJ (Schindelin et al.,2012). Nuclear and cytoplasmic intensity ratio: Regions of interested were manually drawn on respective tissues. Nuclear to cytoplasmic intensity ratio was then calculated using a Fiji/ImageJ macro developed by Volker Baecker at Montpellier RIO Imaging (Intensity Ratio Nuclei Cytoplasm Tool, [RRID:SCR_018673](https://scicrunch.org/scicrunch/Resources/record/nlx_144509-1/SCR_018573/resolver?q=intensity%20ratio%20nuclei%20cytoplasm&l=intensity%20ratio%20nuclei%20cytoplasm)). Documentation can be accessed [here](https://github.com/MontpellierRessourcesImagerie/imagej_macros_and_scripts/wiki/Intensity-Ratio-Nuclei-Cytoplasm-Tool). Same analysis parameters were used across samples compared.

#### Electron micrographs

For morphological sections, composite images were assembled in Fiji (Schindelin et al., 2012) using ImageJ TrackEM2 plugin (Cardona et al., 2012). For immunogold labelling, regions of interest were visually determined and gold particles were hand-counted in Fiji/ImageJ.

#### Wing area

Each wing blade was hand segmented and area measured in pixels using a self-written macro on Fiji/ImageJ.

#### Pupal volume

Using a self-written Fiji/Imagej macro, an ellipse was fitted to the pupa. Volume was calculated assuming an ellipsoid using the formula below (minor axis length as the height) -

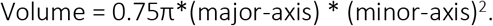

### Data representation and statistical analysis

Data wrangling was done using Microsoft Excel and R-Studio (version 1.3.1093). R-Studio was also used for plotting. Packages used in R- plyr, dplyr, ggplot2 and xlsx. Plots generated through R were then formatted using Adobe Illustrator 2021. Illustrations were generated using BioRender (www.biorender.com). Statistical analysis was done using Microsoft Excel, GraphPad Prism 5.0 and online through https://www.socscistatistics.com. Information about sample sizes and statistical tests used in this study can be found in the respective figure legends. Detailed information on sample size and replicates have also been summarized in Table 1 of the supplementary material.

