## Supplemental Information for "A local insulin reservoir ensures developmental progression in condition of nutrient shortage in *Drosophila*"

Supplementary Table 1

| Figure ref. | n indicates | Values shown on Figures |
|---|---|---|
| 1g | Number of ring glands/larvae used in 1 experiment- 3(control) and 5(Dilp2 RNAi) | Individual CC soma – 12 (control) and 14 (Dilp2 RNAi) |
| 1h | Number of ring glands/larvae used in 1 experiment- 4(control) and 7(Dilp5 RNAi) | Individual CC soma – 30 (control) and 32 (Dilp5 RNAi) |
| 2c | Number of ring glands/larvae used in 1 experiment- 3(control), 4(IMPL2 RNAi) and 5 (InR RNAi) | Individual CC soma – 16 (control) ,23 (IMPL2 RNAi) and 23 (InR RNAi) |
| 2e | Number of ring glands/larvae used in 1 experiment- 3(control) and 4(IMPL2 RNAi) | Individual CC soma – 18 (control) and 15 (IMPL2 RNAi) |
| 3a | Total number of vials (15-25 larvae each) used in 2 independent experiments | 6 (both) |
| 3b | Total number of pupae used in 3 independent experiments | 24 (both) |
| 3c | Total number of vials (8-15 flies each) used in 4 independent experiments | 7 (control) and 9 (IMPL2 RNAi) |
| 3d | Total number of adult flies used in 3 independent experiments | 55 (control) and 20 (IMPL2 RNAi) |
| 3e | Total number of weight measurements (13-15 larvae each) from 2 (96 hrs) or 3 (104 and 116hrs) independent experiments | 8 (96 hrs) and 12 (104 and 116 hrs) |
| 3f | Total number of ELISA assays (pooled from 5-8 larvae) from 2 independent experiments | 7 (control) and 5 (IMPL2 RNAi) |
| 3h | Number of fat bodies (one per larva) from 2 independent experiments | 10 (both) |
| 4d | Number of different microscopic fields used from 1 tissue section | 4 |
| 5b,c | Total number of vials (15-25 larvae each) used in 2 independent experiments | 6 (all) |
| 5e | Number of ring glands/larvae used in 1 experiment- 4 (starved) and 3(fed) | Individual CC soma – 16 (fed) and 17 (starved) |

| | | |
|---|---|---|
| 5g | Number of ring glands/larvae used in 1 experiment- 3 (starved and fed) | Individual CC soma – 14 (fed) and 11 (starved) |
| 5h,i | Total number of ELISA assays (5-8 larvae used for each) from 1 experiment | 5 (fed) and 4 (starved) |
| 5k | Number of fat bodies (one per larva) from 1 experiment | 5 (both) |
| 5m | Number of prothoracic glands (one per larva) from 1 experiment (104 hours) and 3 independent experiments (116 hours) | 4 (104 hours) and 9 (116 hours) |
| S2b | Number of ring glands/larvae used in 1 experiment- 1(both) | Individual CC soma – 7(control) and 8(knock-out) |
| S3 | Number of ring glands/larvae used in 1 experiment- 3 (control), 4 (IMPL2 RNAi) and 4 (InR RNAi) | Individual CC soma – 14 (control) , 14 (IMPL2 RNAi) and 18 (InR RNAi) |

Supplementary Figures

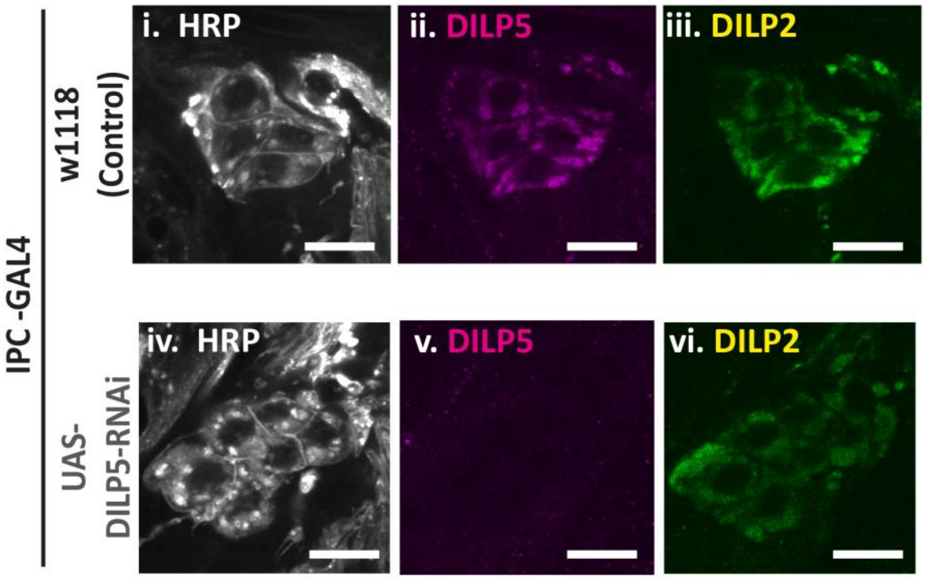


**Figure S1. Corpora cardiaca DILP2 is unaffected in IPC-specific DILP5 knock-down.**

CC soma from IPC-specific(*dilp2*-gal4) DILP5 RNAi (iv-vi) together with wild-type controls(i-iii). Stained for neuronal marker HRP (i and iv, gray), DILP5 (ii and v, magenta) DILP2 (iii and vi, green). Scale bar = 10um.

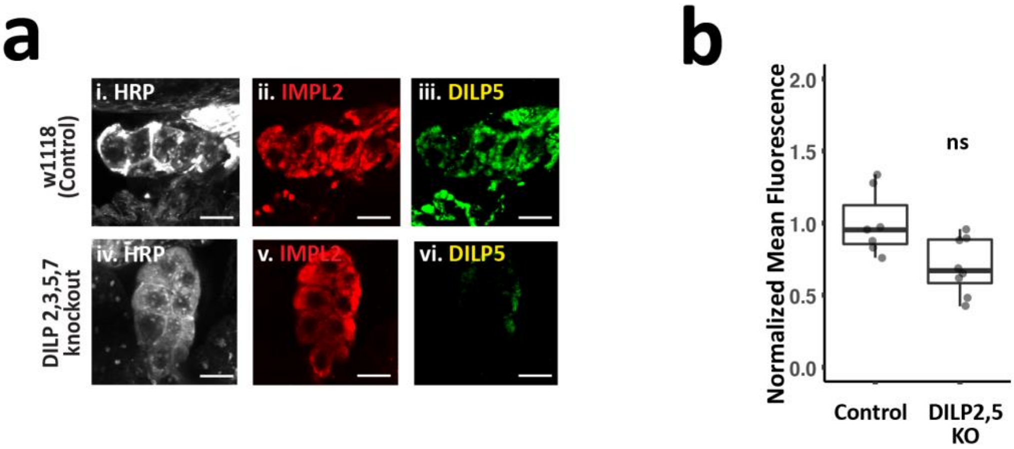


Figure S2. Corpora cardiaca IMPL2 production is independent of DILP2 and 5.

**(a)** CC soma of controls and DILP2,3,5,7 knock-out larvae stained for HRP (i and iv, gray), IMPL2(a.ii and a.iv, red) and DILP5(a.iii and a.vi, green). Scale bar = 10um.

**(b)** Average fluorescence intensity of IMPL2 normalized to the control mean. Each point represents a single CC soma. n =1; ns = $p>0.01$; Student's t-test.

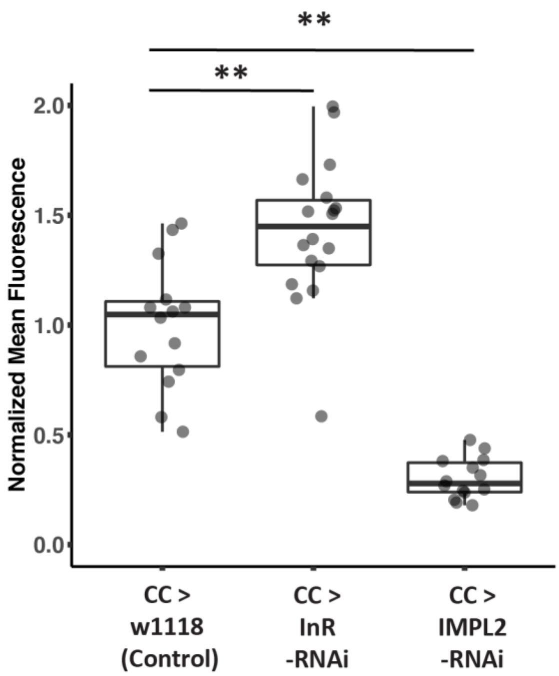


Figure S3. Corpora cardiaca IMPL2 is necessary for DILP5 uptake.

Average fluorescence intensity of DILP5 normalized to the control mean. Each point represents a single CC soma. n = 3-4; **= $p<0.01$; Student's t-test.

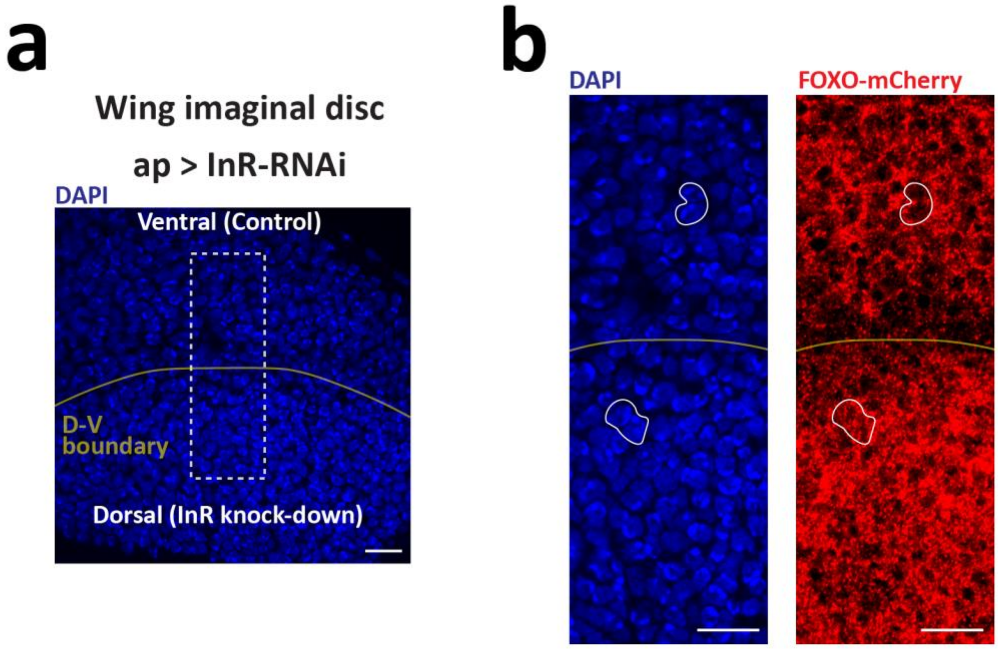


Figure S4. Efficacy of insulin receptor RNAi.

**(a)** Wing pouch nuclei (DAPI, blue) of in L3 wing imaginal disc segmented into dorsal and ventral compartment. Insulin receptor RNAi expression only in the dorsal compartment (*ap*-gal4, tub-gal80TS) for 24hours at 29˚C . Dashed white area magnified in **(b).**

**(b)** Also Stained for mCherry(red) in larvae containing transgene FOXO-mCherry. Solid white outlines show a nuclei in dorsal or ventral compartment. Nuclear localization of FOXO-mCherry indicate insulin pathway activity. Dorso-ventral (D-V) boundary is denoted by yellow line. Scale = 10um.

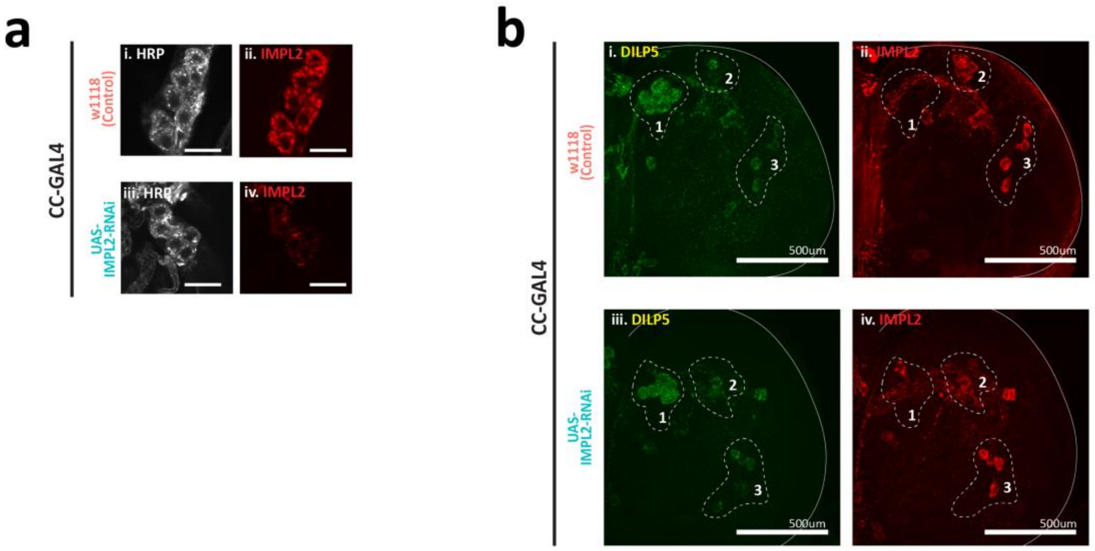


**Figure S5. Efficacy of IMPL2 RNAi.**

CC-specific(*akh*-gal4) RNAi-mediated knockdown of IMPL2 and wild-type control.

**(a)** CC soma stained for neuronal marker HRP (a.i and a.iii, gray) and IMPL2 (a.ii and a.iv, red). Scale = 10um.

**(b)** z-projection of the right hemisphere of L3 larval brain, stained for DILP5 (b.i and b.iii, green) and IMPL2 (b.ii and b.iv, red). Numbered outlines indicate (1) insulin-producing cells (IPCs), (2) and (3) IMPL2- producing neurons. Scale bar = 500um.
